# Virome analysis of New Zealand’s bats reveals cross-species viral transmission among the *Coronaviridae*

**DOI:** 10.1101/2023.06.19.545647

**Authors:** Stephanie J. Waller, Pablo Tortosa, Tertia Thurley, Colin O’Donnell, Rebecca Jackson, Gillian Dennis, Rebecca M. Grimwood, Edward C. Holmes, Kate McInnes, Jemma L. Geoghegan

## Abstract

The lesser short-tailed bat (*Mystacina tuberculata*) and the long-tailed bat (*Chalinolobus tuberculatus*) are Aotearoa New Zealand’s only native extant terrestrial mammals and are believed to have migrated from Australia. Long-tailed bats arrived in New Zealand an estimated two million years ago and are closely related to other Australian bat species. Lesser short-tailed bats, in contrast, are the only extant species within the Mystacinidae and are estimated to have been living in isolation in New Zealand for the past 16-18 million years. Throughout this period of isolation, lesser short-tailed bats have become one of the most terrestrial bats in the world.

Through a metatranscriptomic analysis of guano samples from eight locations across New Zealand we aimed to characterise the viromes of New Zealand’s bats and determine whether viruses have jumped between these species over the past two million years. High viral richness was observed among long-tailed bats with viruses spanning seven different viral families. In contrast, no bat-specific viruses were identified in lesser short-tailed bats. Both bat species harboured an abundance of likely dietary– and environmental-associated viruses. We also identified alphacoronaviruses in long-tailed bat guano that had previously been identified in lesser short-tailed bats, suggesting that these viruses had jumped the species barrier after long-tailed bats migrated to New Zealand. Of note, an alphacoronavirus species discovered here possessed a complete genome of only 22,416 nucleotides with entire deletions or truncations of several non-structural proteins, thereby representing what is possibly the shortest genome within the *Coronaviridae* identified to date. Overall, this study has revealed a diverse range of novel viruses harboured by New Zealand’s only native terrestrial mammals, in turn expanding our understanding of bat viral dynamics and evolution globally.

## 2. Introduction

Bats harbour a wide diversity of viruses, some of which have the potential to spillover to other hosts, including humans (1,2). Viruses such as lyssaviruses, coronaviruses, Ebola, Hendra and Nipah are all carried by bats, with significant public health and economic consequences (3–10). Accordingly, bats have become the focus of many virological surveys to help identify novel viruses and assess the risk they pose for zoonoses (11). Despite this, investigations into the viruses present in the native bat species of New Zealand have been limited.

New Zealand is home to only two native extant terrestrial mammals; the lesser short-tailed bat (*Mystacina tuberculata*) and the long-tailed bat (*Chalinolobus tuberculatus*) (12). A third species, the greater short-tailed bat, *Mystacina robusta*, has not been sighted since 1967 and is likely extinct (12). The lesser short-tailed bat can be further classified into three subspecies: the northern lesser short-tailed bat (*Mystacina tuberculata aupourica*), the central lesser short-tailed bat (*Mystacina tuberculata rhyacobia*) and the southern lesser short-tailed bat (*Mystacina tuberculata tuberculata*). An additional two bat species have been identified in the New Zealand fossil record; *Vulcanops jennyworthyae* and *Mystacina miocenalis* (13,14). Viral sequences from the *Papillomaviridae*, *Polymaviridae*, *Caliciviridae*, *Hepeviridae*, *Poxviridae*, *Parvoviridae*, *Adenoviridae*, *Picornaviridae* and *Coronaviridae* families have previously been uncovered in lesser short-tailed bats located on a remote, predator-free, offshore island (15,16), while viral transcripts from the *Coronaviridae* have previously been uncovered in mainland long-tailed bats and lesser short-tailed bats using a pan-coronavirus PCR approach (17). Building on this previous work, we aimed to use total RNA sequencing to document complete viromes in these species and, from this, determine whether viruses have exhibited cross-species transmission between New Zealand’s bat species.

Bats are believed to be an important reservoir of a multitude of viruses in part because they roost in large numbers, exhibit species co-habitation and are able to fly, readily transferring viruses among geographic regions. In addition, bats are thought to be able to tolerate a high abundance of viruses due to unique components of their immune system (18–20). As of 2021, viruses spanning a total of 24 RNA viral families and 11 DNA viral families have been uncovered in hosts belonging to the order Chiroptera (18). Of the bat-associated viral sequences published in GenBank, 85% were RNA viruses, of which 30% and 24% were from the families *Coronaviridae* and *Rhabdoviridae*, respectively (18).

Around 84 million years ago the continental crust from which New Zealand was formed separated from Gondwana (21). Since mammals evolved ∼32-27 million years ago, all terrestrial mammals present in New Zealand originated from overseas with their arrival aided by winds, ocean currents and human movement (12). The long-tailed bat is believed to have arrived from Australia during the Pleistocene around two million years ago (12). It is a member of the Vespertilionidae family, diverging from the last common ancestor approximately 17 million years ago, such that it could have migrated to New Zealand earlier than previously thought (22). The Vespertilionidae, that comprises of more than 360 species, is considered one of the most widely dispersed mammalian families in the world (12). In comparison, the lesser short-tailed bat is the sole surviving member of the Mystacinidae and based on fossil records migrated from Australia during the early Miocene period approximately 16-18 million years ago (23,24). Around this time, the remainder of the mystacinid lineage became extinct in Australia (12). Mitochondrial phylogenetic analysis indicates that the lesser short-tailed bat last shared a common ancestor with the Noctilionoidae superfamily around 35 million years ago (25), again suggesting this species may have arrived earlier than fossil records propose.

The viral diversity of two species of bats from Australia, Gould’s wattled bat (*Chalinolobus gouldii*) and chocolate wattled bat (*Chalinolobus morio*), which are closely related to New Zealand’s long-tailed bats, has previously been analysed. Viruses from the *Coronaviridae*, *Adenoviridae* and *Paramyxoviridae* families have previously been identified in both these species (26). Of note, the coronaviruses found in Gould’s wattled bats were closely related to the coronaviruses previously detected in New Zealand’s long-tailed bats, suggesting that virus-host codivergence over extended time scales has likely played a role in their evolution (17,26).

Due to its isolation in New Zealand, the lesser short-tailed bat has developed several atypical behaviours distinct from any other known bat species. In particular, the lesser short-tailed bat is considered the most terrestrial bat in the world, such that it has adapted to spending large periods of time on the forest floor foraging for food (12,27). Additionally, the diet of lesser short-tailed bats is considered one of the broadest of any bat species, including nectar, fruit, pollen, terrestrial and flying invertebrates, as well as potentially fungi. This diet allows them to remain active for most of the year unlike most other non-tropical insectivore bat species (12,27,28). In contrast, long-tailed bats are strictly insectivores and spend more time flying in comparison to lesser short-tailed bats (12,28). During winter, long-tailed bats enter a state of short-term torpor as the number of flying insects, their main food source, reduces (12,29).

Herein, we sampled guano from both long-tailed and lesser short-tailed bats. We investigated the viromes of these species to determine if viruses have jumped between these hosts and test whether the marked differences in their ecology and behaviour shaped the composition of their virome.

## 3. Materials and Methods

### 3.1. Bat Guano Sample Collection

Guano from lesser short-tailed bats and long-tailed bats was collected as part of several annual monitoring programmes carried out by the New Zealand Department of Conservation. Guano was collected using tarps, harp traps and catch bags from eight locations/roosts across New Zealand (Figure 1). A total of 219 individual guano samples were collected during two sampling periods: February 2020 and January-February 2021. More information regarding sample locations, species, as well as the number of individual guano samples within each group, is provided in Supplementary Table 1. Guano samples were submerged into 1mL of RNAlater and briefly stored at 4 °C until they were sent to the University of Otago, Dunedin, where they were stored at –70 °C or –80°C until total RNA was extracted.

**Figure 1.**
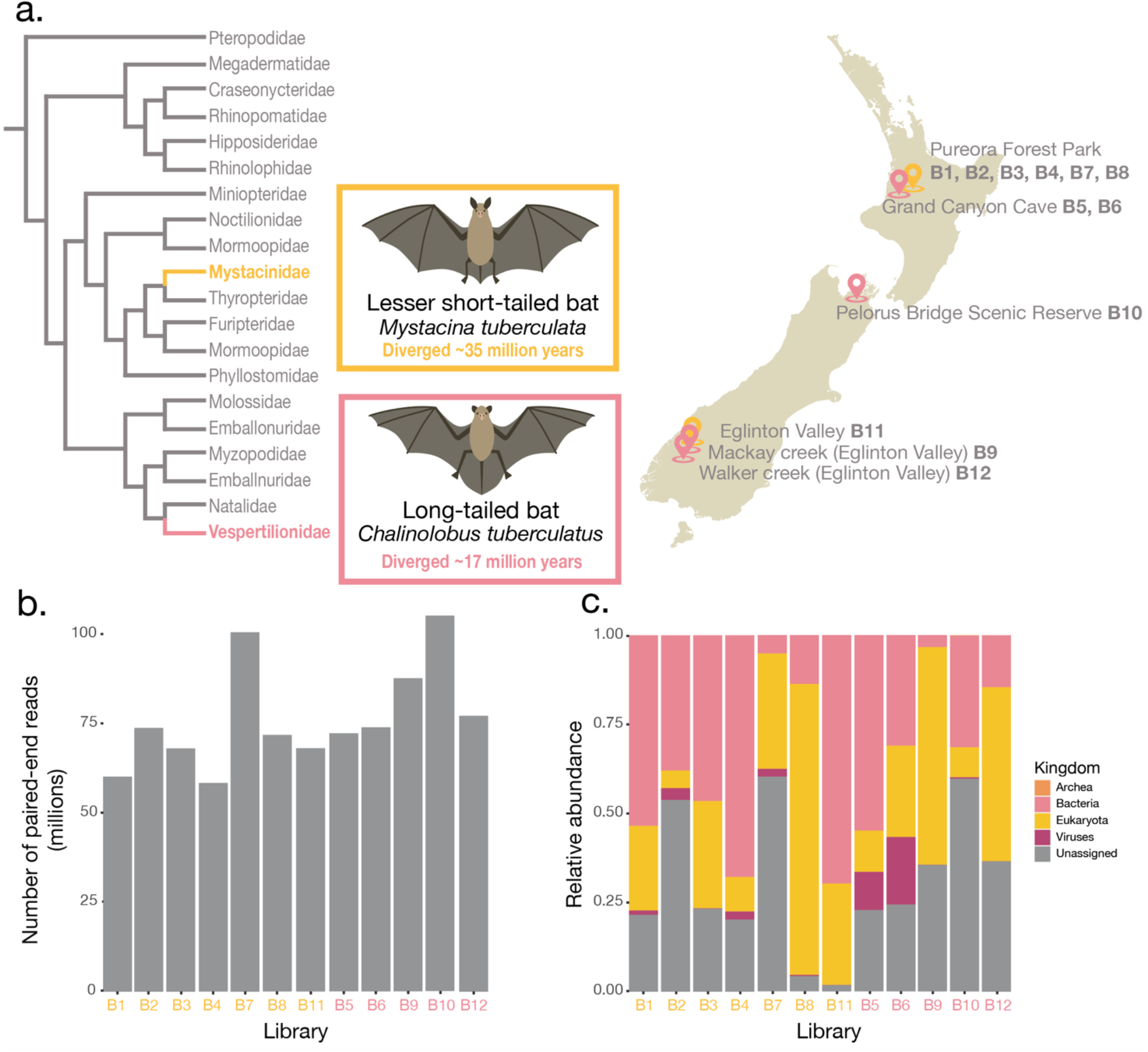
(**a**) A cladogram (left) illustrating the evolutionary relationships of bat families within the Chiroptera order. The *Mystacina tuberculata* species is a part of the Mystacinidae, highlighted in yellow which diverged from other bats ∼35million years ago (25,47) while the Vespertilionidae, highlighted in pink, includes the *Chalinolobus tuberculatus* species which is believed to have diverged from other species within the Vespertilionidae ∼17 million years ago (22). A map of New Zealand (right) indicating the sampling locations and roosts of pooled bat guano samples. Bat illustrations were provided by Hamish Thompson and were used with permission. (**b**) Total paired– end sequencing reads from bat guano metatranscriptome libraries. (**c**) Relative abundance of the standardised abundances of archea, bacteria, eukaryota, viruses and unassigned contigs, based on the BLASTn sskingdom function. The total estimated abundances across each library were first standardised by the total number of reads within each library before being normalised by the total standardised abundance within a given library.

### 3.2. Bat Guano Total RNA Extraction

Frozen bat guano samples stored in RNAlater were first defrosted. Approximately 25-50mg of the defrosted guano was placed in 15mL RNase-free round bottom tubes. In a fume hood, 500ul of cold TRIzol^TM^ was added and samples were homogenised for one minute using a TissueRuptor (Qigaen). The homogenate was then centrifuged at 4°C at 12,000 xg for five minutes. Total RNA from the clear supernatant was then extracted following the TRIzol manufacturers protocol with minor alterations. Briefly, a secondary chloroform phase separation step was added to remove any residual phenol contamination while an additional ethanol wash step was added to remove any residual guanidine contamination. Extracted RNA was quantified using a Nanodrop. RNA from 217 of the 219 guano samples was at suitable concentrations for downstream processing. Equal concentrations of RNA from 5-41 individuals were pooled into 12 groups based on bat species and sample location (see Supplementary Table 1).

### 3.3. Bat Guano RNA Sequencing

Extracted RNA was subject to total RNA sequencing. Libraries were prepared using the Illumina Stranded Total RNA Prep with Ribo-Zero Plus (Illumina) and 16 cycles of PCR. Paired-end 150bp sequencing of the RNA libraries was performed on the Illumina NovaSeq 6000 platform using a single S4 lane.

### 3.4. Virome Assembly and Virus Identification

Paired reads were trimmed and assembled *de novo* using Trinity v2.11 with the “trimmomatic” flag option (30). Sequence similarity searches against the NCBI nucleotide (nt) database (2021) and the non-redundant (nr) protein database (2021) using BLASTn and Diamond (BLASTx), respectively, were used to annotate assembled contigs (31). Contigs were categorised into higher kingdoms using the BLASTn “sskingdoms” flag option. Non-viral blast hits including host contigs with sequence similarity to viral transcripts (e.g. endogenous viral elements) were removed from further analysis during manual screening. A maximum expected value of 1×10^-10^ was used as a cut-off to filter putative viral contigs. Viral contigs that have previously been identified as viral contaminants from laboratory components were also removed from further analysis (32). Based on the BLASTn and Diamond results (database accessed November 2022), putative viral contigs were further analysed using Geneious Prime 2022.2.2 to find and translate open reading frames (ORFs).

### 3.5. Protein Structure Similarity Searching for Viral Discovery

Similar to Waller et al. 2022 (33), we used a protein structure similarity search to identify highly divergent viral transcripts that did not share significant amino acid sequence similarity to other known transcripts. Such “orphan contigs” (34) were translated into ORFs using the EMBOSS getorf program (35), with the minimum nucleotide size of the ORF set to 1,000 nucleotides, the maximum nucleotide size of the ORF set to 50,000 and the “methionine” flag option set to only report ORFs with a methionine amino acid starting codon. Reported ORFs were submitted to Phyre2, which uses remote homology detection to build 3D models to predict and analyse protein structure and function (36). Virus sequences with predicted polymerase structures with a confidence value of >90% were aligned with representative amino acid sequences from the same viral family obtained from NCBI RefSeq using MAFFT v7.490 (L-INS-I algorithm). Conserved domains were visually confirmed before phylogenetic trees were estimated using the same method outlined in section 3.7.

### 3.6. Estimating Viral Transcript Abundance Estimations

Viral abundances were estimated using Trinity’s “align and estimate” tool. RNA-Seq by Expectation-Maximization (RSEM) (37) was selected as the method of abundance estimation, Bowtie2 (38) as the alignment method and the “prep reference” flag enabled. To mitigate the impact of contamination due to index-hopping, viral transcripts with expected abundances of less than 0.1% of the highest expected abundance for that virus across other libraries were removed from further analysis. Total viral abundance estimates for viruses from vertebrate hosts (i.e. bats), vertebrate-associated hosts (i.e. where the true host of the virus remained unclear) or viruses from both vertebrate hosts and vertebrate-associated hosts across viral families and orders were compiled across all libraries. Estimated abundances were standardised to the number of paired reads per library.

### 3.7. Virus Phylogenetic Analysis

Translated viral protein polymerase sequences (i.e. RNA-dependent RNA polymerase, RdRp) or capsid sequences were aligned with representative protein sequences from the same taxonomic viral family or order obtained from NCBI RefSeq as well as the closest BLAST hit using MAFFT v7.450 (L-INS-I algorithm) (see Supplementary Table 2 for lengths of sequence alignments) (39). Poorly aligned regions were removed using trimAL v1.2rev59 with the gap threshold flag set to 0.9 (40). IQ-TREE v1.6.12 was used to estimate maximum likelihood phylogenetic trees for each viral species/family/order (41). The LG amino acid substitution model was selected with 1000 ultra-fast bootstrapping replicates and the –alrt flag specifying 1000 bootstrap replicates for the SH-like approximate likelihood ratio test. Phylogenetic trees were annotated using Figtree v1.4.4 (42).

### 3.8. Analysis of Alpha Diversity on Virome Composition

All statistical analysis plots were created using RStudio v2021.09.2 with the tidyverse ggplot2 package (43). Using the diversity analysis function which is part of the vegan package (44), the Shannon index was selected as the index method to measure alpha diversity of standardised abundance estimates of viral families/orders across bat species and location (North Island or South Island). A Welches T-test was used to determine whether there was a significant difference in virome alpha diversity across bat species and location (North Island and South Island). The cor and cor.test function in R was used to test the correlations between the pairwise differences in Shannon Index and the pairwise distance between roost site (km).

### 3.9. Viral Nomenclature

A virus was arbitrarily considered a novel species if it shared <90% amino acid similarity within the most conserved region (i.e. RdRp/polymerase or capsid) (45,46), unless otherwise stated. For novel virus sequences we have provided a proposed virus name (prior to formal verification by the International Committee on Taxonomy of Viruses (ICTV)). Viruses that were most closely related to a virus identified in a vertebrate metagenome sample, such that their true host was uncertain, as well as invertebrate host-associated viruses, were classified as ‘bat metagenome derived’ viruses as these viruses were unlikely to directly infect bats. We have also proposed the general name, pekapeka where viruses were found in both bat species (Māori refer to New Zealand’s bats generally as pekapeka).

## 4. Results

Total RNA isolated from 219 bat guano samples were pooled into 12 libraries based on bat species, sampling location/roost and sampling year and were subject to whole RNA sequencing (Supplementary Table 1). The number of sequencing reads generated from the 12 bat guano pools varied between 58-105 million paired-end reads per library (Figure 1b, Supplementary Table 1). Viral contigs contributed 0.02-18.9% of the total standardised abundances across each of the bat guano pooled samples, while contigs that could not be assigned to a source based on sequence similarity (i.e. orphan contigs) made up 1.75-60.24% of the total standardised abundances (Figure 1c).

### 4.1 Viral Abundance and Diversity

Analysis of bat guano samples revealed that the viromes of long-tailed bats were more diverse than those of lesser short-tailed bats, with samples of the former containing bat viral contigs spanning seven viral families (*Picornaviridae*, *Astroviridae*, *Paramyxoviridae*, *Papillomaviridae*, *Poxviridae*, *Rhabdoviridae* and *Coronaviridae*) (Figure 2). Surprisingly, no bat-specific viral contigs were identified in the lesser short-tailed bat samples in this study, although short transcripts of bat papillomaviruses, coronaviruses, hepeviruses, polyomaviruses and caliciviruses have previously been uncovered in lesser short-tailed bats in New Zealand using Illumina MiSeq2000 and PCR approaches (16,17). Both bat species examined here contained bat ‘metagenome derived’ viral contigs (Figure 2). These represent viral contigs in which the true host is uncertain but were likely dietary– or environmental-associated viruses. Long-tailed bat samples contained metagenome derived viral contigs spanning the *Caliciviridae* and *Adenoviridae* families, and data from both bat species contained metagenome derived viral contigs from Basavirus sp., *Flaviviridae*, *Hepeviridae*, *Nodaviridae*, *Peribunyaviridae*, *Picobirnavirdae* and *Reovirales* (Figure 2).

**Figure 2.**
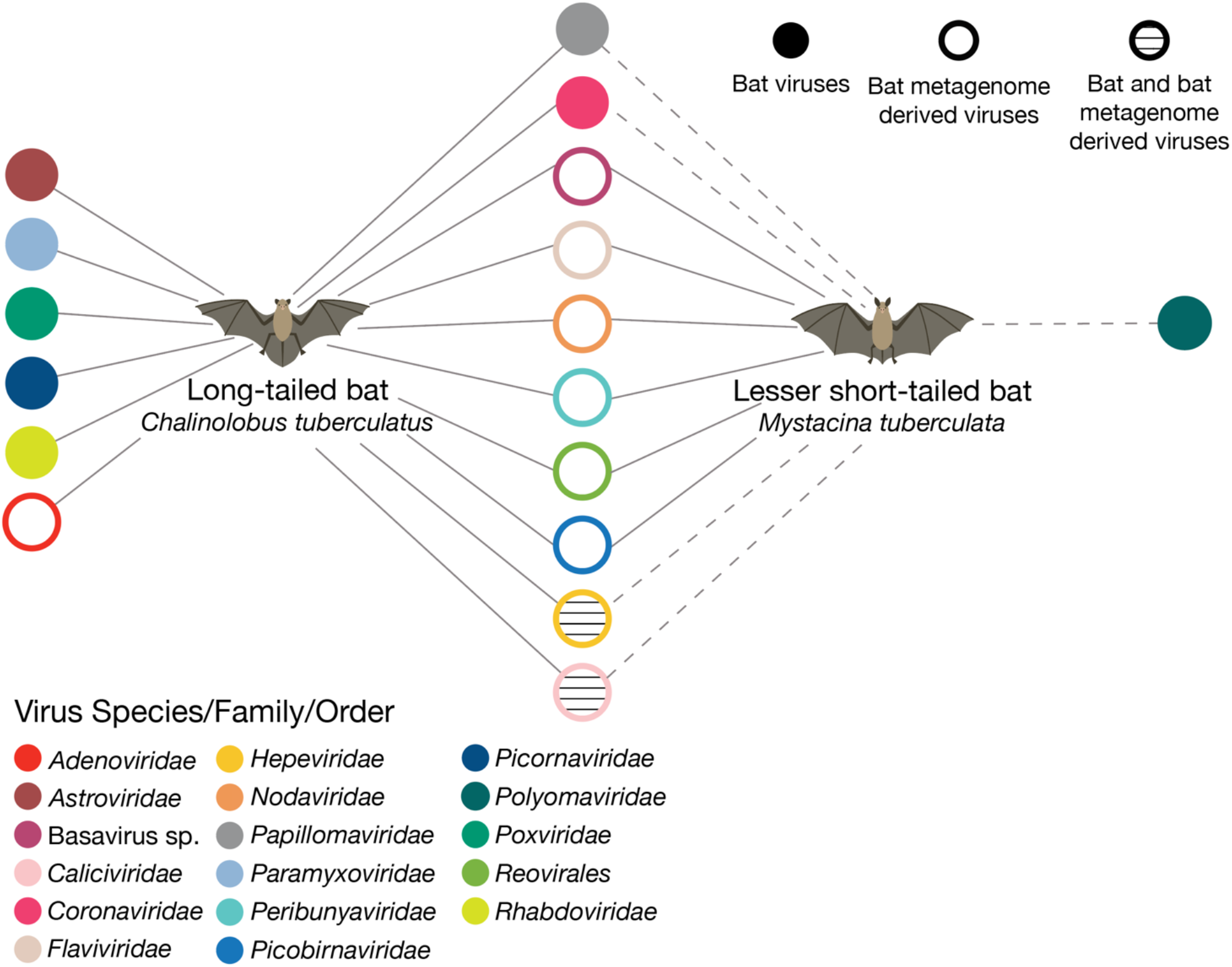
Bipartite network depicting viral families detected in guano samples from long-tailed and lesser short-tailed bats. Where viral families are shared between both long-tailed and lesser short-tailed bats the networks between the two bat species are connected. The dashed line indicates viral transcripts identified in previous studies in lesser short-tailed bats (15–17). Filled circles represent bat viral contigs for a given viral species/family/library, whereas open circles represent bat metagenome derived viral contigs where the true host of the viruses remains unclear.

### 4.2 Novel Bat Viruses

#### 4.2.1 Coronaviridae

We identified several alphacoronavirus full genomes from long-tailed bat guano sampled from the Grand Canyon Cave (Figure 3a) and identified two species of alphacoronaviruses. One alphacoronavirus ORF1b transcript, which contains the RdRp, identified here shared 90.22% amino acid similarity with *Alphacoronavirus sp.* from Gould’s wattled bats from Australia, suggesting that the same virus species infects hosts from Australia and New Zealand based on ICTV species definitions (Figure 3b, Supplementary Table 2) (26,46). Phylogenetic analysis of this region revealed that this transcript was also likely the same viral species as two previously identified viruses, *Mystacina coronavirus New Zealand/2013* and *Chalinolobus tuberculatus alphacoronavirus*, although these previously published viral sequences were very short in length (186 and 129 amino acids, respectively) and hence were not a significant match due to inadequate (∼10%) query coverage. The viral transcript identified here shared amino acid sequence similarity of 98% and 100% with *Mystacina coronavirus New Zealand/2013* and *Chalinolobus tuberculatus alphacoronavirus*, respectively. The former virus was previously uncovered in lesser short-tailed bats sampled from the remote, offshore New Zealand island, Whenua Hou in 2013 (15), while the latter was uncovered and classified in long-tailed bats from the Grand Canyon Cave and sampled in 2020 (Figure 3a) (17).

**Figure 3.**
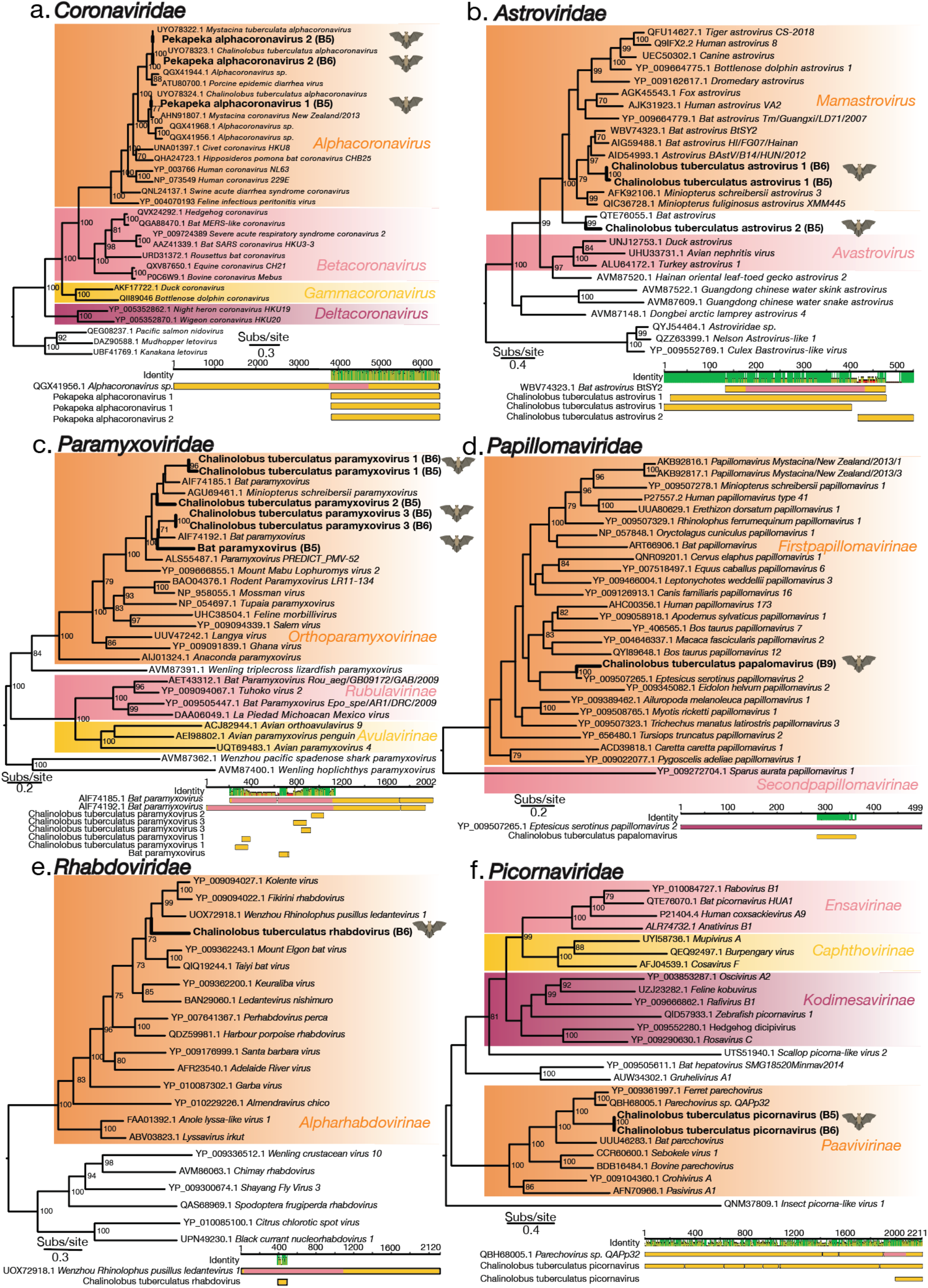
Maximum likelihood phylogenetic trees of representative viral transcripts containing the RdRp from the families (**a**) *Astroviridae*, (**b**) *Coronaviridae*, (**c**) *Paramyxoviridae*, (**e**) *Rhabdoviridae* and (**f**) *Picornaviridae* and (**d**) the major capsid protein from the *Papillomaviridae*. Bat viruses identified in this study are in bold while known genera and subfamilies are highlighted. Branches are scaled to the number of amino acid substitutions per site. All phylogenetic trees, with the exception of that of the *Coronaviridae* which was manually rooted, were midpoint rooted. Nodes with ultrafast bootstrap values of >70% are noted. Below each phylogenetic tree is the genomic organisation that shows the closest BLASTp hit, with the polymerase or capsid region highlighted in pink or dark pink, respectively. Novel bat viral contigs have been aligned to the closest BLASTp hits and the mean pairwise identity of the alignment (green bars) as well as the amino acid length of the alignment are shown.

A full genome was assembled for this alphacoronavirus species revealed that this genome was markedly short in length, comprising only 22,416 nucleotides (Figure 4). Upon further investigation of its genome organisation, the short coronavirus, which was relatively highly abundant with a standardised abundance of 6.5×10^-4^, did not encode the non-structural proteins (NSP) 1 and 2, while NSP3 appeared to be truncated in comparison to other bat alphacoronaviruses (Figure 4). Additionally, while ORF1b shared >90% amino acid similarity with ORF1ab from *Alphacoronavirus sp.* (MK472068.1) (26), other ORFs shared only 32% – 65% amino acid sequence similarity. Given this large deletion in ORF1a, we suggest that this coronavirus be classified as a new species, Pekapeka alphacoronavirus 1.

**Figure 4.**
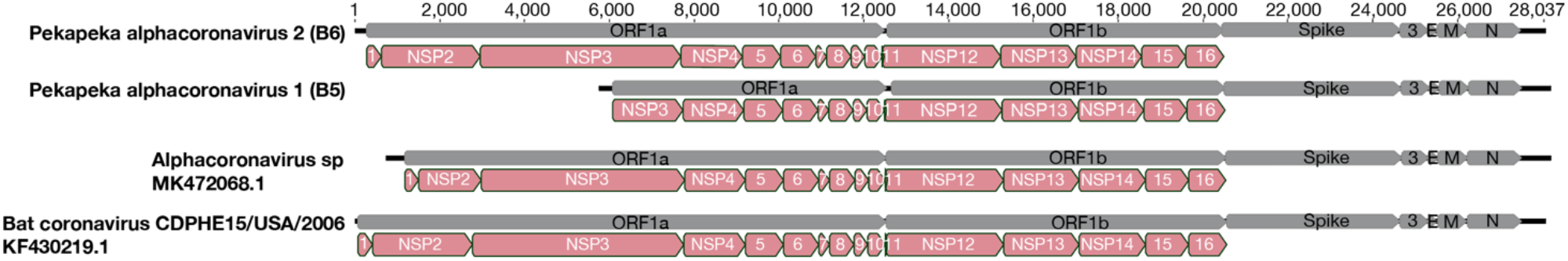
Genome organisation of bat alphacoronaviruses. Bat coronavirus CDPHE15/USA/2006 (KF430219.1) had previously been annotated to depict the positions of the mature peptides within the ORF1a and ORF1b regions. The annotated Bat coronavirus CDPHE15/USA/2006 (KF430219.1) was used as a guide to indicate the position of the mature peptides within Alphacoronavirus sp. isolate WAAlc1 (MK472071.1), Pekapeka alphacoronavirus 2 (B6) and Pekapeka alphacoronavirus 1 (B5).

The second alphacoronavirus species we identified shared 87% amino acid sequence similarity with the closest known genetic relative, *Alphacoronavirus sp.,* again identified in Gould’s wattled bat in Australia (26) (Figure 3a, Supplementary Table 2). Phylogenetic analysis showed that two previously identified viruses, *Chalinolobus tuberculatus alphacoronavirus* in long-tailed bats (17) and *Mystacina tuberculata alphacoronavirus* in lesser short-tailed bats (17), were likely the same viral species. These previously identified virus sequences were short in length (133 amino acids) compared to the full viral genome identified here. While these short sequences shared 100% amino acid identity to the virus we identified here, the query coverage of just 5% meant they were not considered significant matches. Nevertheless, assuming these are the same viral species, we propose the revised name, Pekapeka alphacoronavirus 2, reflecting that this virus infects both New Zealand’s bat species.

#### 4.2.2 Astroviridae

Three novel viral polymerase contigs were identified in long-tailed bat guano sampled from the Grand Canyon Cave that shared sequence similarity to viruses within the *Astroviridae* (positive–sense, single stranded (+ssRNA)) (Figure 3b). Two of these contigs shared >90% amino acid sequence similarity with each other and spanned a similar region of the polymerase protein, indicating that these contigs were a singular viral species, tentatively named Chalinolobus tuberculatus astrovirus 1 (Supplementary Table 2). Chalinolobus tuberculatus astrovirus 1 contigs fell within the genus *Mamastroviruses* (Figure 3b) and shared 71% amino acid sequence similarity with the closest known genetic relative, *Bat astrovirus BtSY2*, which was identified in pomona roundleaf bat (*Hipposideros pomona*) in China (48). The third astrovirus contig shared 78.3% amino acid similarity with the closest known genetic relative, *Bat astrovirus*, previously identified in a common vampire bat (*Desmodus rotundus*) in Peru (49).

#### 4.2.3 Paramyxoviridae

Six *Paramyxoviridae* contigs were uncovered in long-tailed bat guano metatranscriptomes sampled from the Grand Canyon Cave. All six *Paramyxoviridae* contigs identified here fell within the subfamily *Orthoparamyxovirinae* (Figure 3c). Two of these contigs shared >90% amino acid sequence similarity with each other and were provisionally named, Chalinolobus tuberculatus paramyxovirus 1 (Figure 3c). Chalinolobus tuberculatus paramyxovirus 1 shared ∼73% amino acid sequence similarity with the closest known genetic relatives, *Bat paramyxovirus* and *Miniopterus schreibersii paramyxovirus*, which were previously uncovered in greater tube-nosed bats (*Murina leucogaster*) and common bent-wing bats (*Miniopterus schreibersii*) from China (Supplementary Table 2) (50).

Another contig that was identified here shared 86% amino acid sequence similarity with the closest known genetic relative, *Bat paramyxovirus*, which was uncovered in Daubenton’s bats (*Myotis daubentonii*) from China (50). This contig was provisionally named Chalinolobus tuberculatus paramyxovirus 2. A further two contigs shared greater than 90% similarity with each other and were consequently named Chalinolobus tuberculatus paramyxovirus 3. This virus shared ∼87% amino acid similarity with the closest known genetic relatives, *Paramyxovirus PREDICT_PMV-52* and *Bat paramyxovirus*, previously uncovered in western bent-winged bats (*Miniopterus magnater*) from Thailand and greater tube-nosed bats from China, respectively (50). The final transcript belonging to the *Paramyxoviridae* we uncovered shared 92% amino acid sequence similarity with *Bat paramyxovirus*, which was previously uncovered in greater tube-nosed bats from China (50). The close genetic match suggests that the same virus species is also present in New Zealand.

#### 4.2.4 Papillomaviridae

A viral transcript sharing amino acid sequence similarity to the *Papillomaviridae* (a double-stranded DNA (dsDNA) virus) was identified in bat guano from long-tailed bats sampled from Mackay Creek in the Eglington Valley. This contig, termed Chalinolobus tuberculatus papillomavirus, shared 88.46% amino acid sequence similarity with the closest known genetic relative, *Eptesicus serotinus papillomavirus 2*, which was identified in serotine bats (*Eptesicus serotinus*) from Spain (Figure 3d, Supplementary Table 2) (51). The novel Chalinolobus tuberculatus papillomavirus along with *Eptesicus serotinus papillomavirus 2* fell within the *Firstpapillomavirinae* subfamily.

#### 4.2.5 Rhabdoviridae

We identified a contig from the *Rhabdoviridae* (negative-sense, single stranded (-ssRNA)) in long-tailed bat guano sampled from Grand Canyon Cave. This viral contig shared 62.63% amino acid sequence similarity with the closest known genetic relative, *Wenzhou Rhinolophus pusillus ledantevirus 1* sampled from least horseshoe bats (*Rhinolophus pusillus*) in China (Supplementary Table 2). Termed Chalinolobus tuberculatus rhabdovirus, this virus fell within the *Alpharabdovirinae* subfamily alongside other species from the *Alpharabdovirinae* identified in Chinese rufous horseshoe bats (*Rhinolophus sinicus*) (52), as well as eloquent horseshoe bats (*Rhinolophus eloquens*) and striped leaf-nosed bats (*Macronycteris vittatus*) from Kenya (Figure 3e) (53,54).

#### 4.2.6 Picornaviridae

Two contigs from the *Picornaviridae* were identified in bat guano from long-tailed bats sampled from the Grand Canyon Cave. Both contigs shared 100% amino acid sequence similarity with each other and were provisionally named Chalinolobus tuberculatus picornavirus. The Chalinolobus tuberculatus picornavirus fell within the *Paavivirnae* subfamily alongside the closest known genetic relatives, *Parechovirus sp. QAPp32* (amino acid sequence similarity 61.21%) identified in the intestinal sample of the common pipistrelle bat (*Pipistrellus pipistrellus*) from China, as well as *Ferret parechovirus* (amino acid sequence similarity 55.46%) identified in a rectal swab from ferrets in the Netherlands (55) (Figure 3f, Supplementary Table 2).

### 4.3 Novel Bat Metagenome Derived Viruses

#### 4.3.1 Basavirus sp

Three Basavirus contigs were identified in bat guano from lesser short-tailed bats sampled from Pureora Forest Park, two of which shared 100% amino acid sequence similarity and were termed Mystacina tuberculata metagenome derived basavirus 1 (Figure 5a, Supplementary Table 2). The Mystacina tuberculata metagenome derived basavirus 1 was most closely related to *Picornavirales sp.*, previously identified in a soil sample from China (56) (amino acid sequence similarity 63%) and *Fisavirus* 1, previously identified in the intestinal contents of a common carp (*Cyprinus carpio*) from Hungary (amino acid sequence similarity 59%) (57). The third contig shared 56% amino acid sequence similarity with the closest genetic relative, *Basavirus sp.*, which was identified in the guano of a lesser Asiatic yellow bat (*Scotophilus kuhlii*) from Vietnam (Figure 5a, Supplementary Table 2) (58). Although these viruses were related to other bat viruses, the true host of these basavirus species is unclear since such divergent members of the *Picornavirales* are often found within the fecal viromes of vertebrates (58). Consequently, we have termed these novel viruses ‘bat metagenome derived’ since there is a strong possibility they are of dietary, environmental, commensal, or parasitic origin.

**Figure 5.**
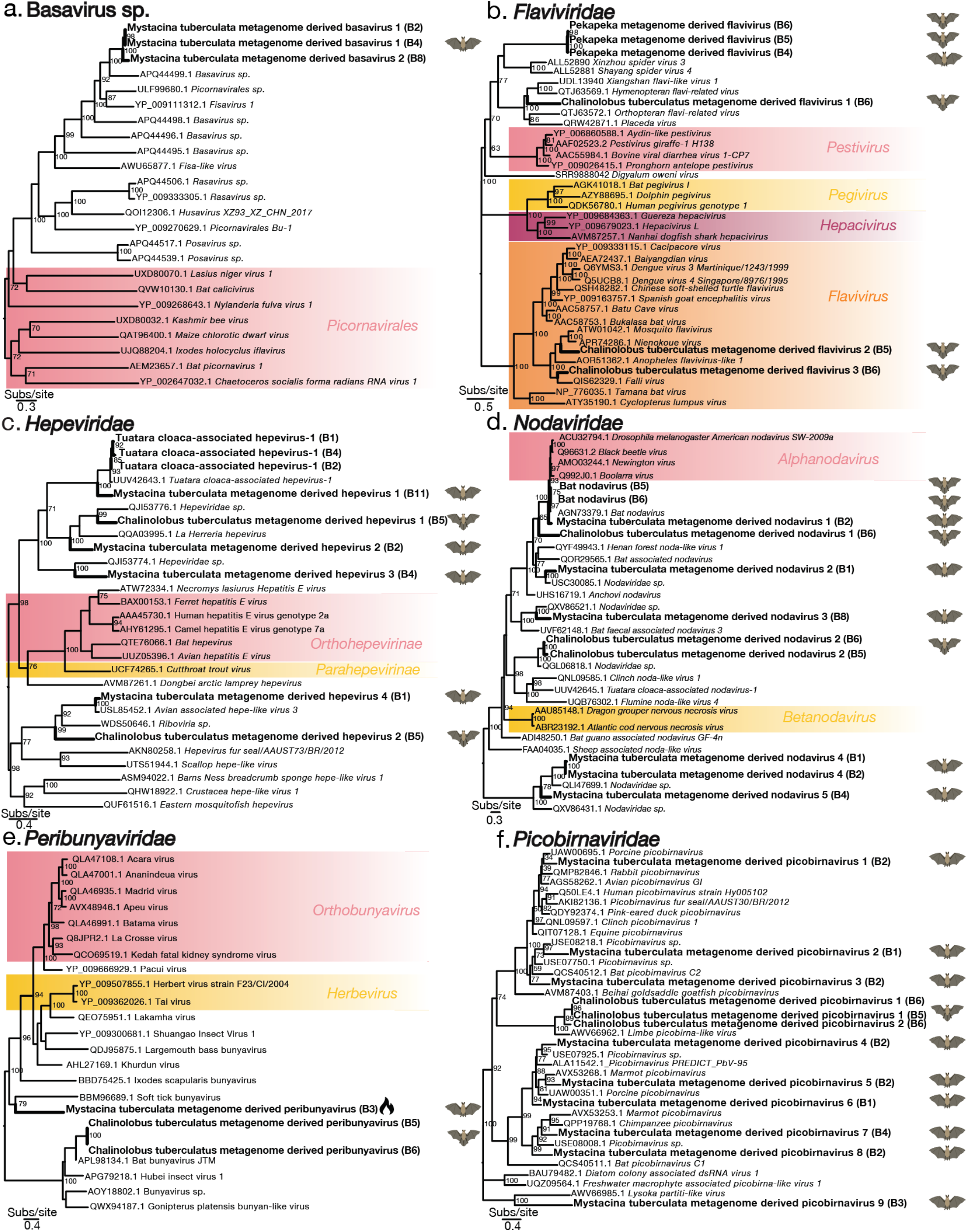
Maximum likelihood phylogenetic trees of representative viral transcripts containing RdRp from the viral species and families (**a**) Basavirus sp., (**b**) *Flaviviridae*, (**c**) *Hepeviridae*, (**d**) *Nodaviridae*, (**e**) *Peribunyaviridae* and (**f**) *Picobirnaviridae*. Bat metagenome derived viruses identified in this study are in bold while viral genera, subfamilies and orders are highlighted. The flame symbol highlights a viral contig that was identified using a protein structural similarity based approach rather than an amino acid sequence similarity based approach. Branches are scaled to the number of amino acid substitutions per site. All phylogenetic trees have been midpoint rooted. Nodes with ultrafast bootstrap values of >70% are noted.

#### 4.3.2 Flaviviridae

We identified four flaviviruses provisionally named, Pekapeka metagenome derived flavivirus, and Chalinolobus tuberculatus metagenome derived flaviviruses 1-3 (Figure 5b, Supplementary Table 2). Viral contigs from Pekapeka metagenome derived flavivirus were identified in both lesser short-tailed and long-tailed bat guano samples (Figure 5b, Supplementary Table 2). As a result, we termed this virus Pekapeka metagenome derived flavivirus reflecting that this virus was identified in both of New Zealand’s bat species. Pekapeka metagenome derived flavivirus fell within the unclassified pesti-large genome flavivirus clade and was highly divergent as indicated by its long branch length on the phylogenetic tree (Figure 5b). The closest BLASTp hit to the Pekapeka metagenome derived flavivirus was to a *Hymenopteran flavi-related virus*, identified in a species of bee (*Thyreus orbatus*) from Italy (amino acid sequence similarity 22-23%) (59) (Supplementary Table 2). Pesti-large genome flaviviruses have previously been associated with invertebrate hosts (60), therefore Pekapeka metagenome derived flavivirus is likely dietary related.

Chalinolobus tuberculatus metagenome derived flavivirus 1 also shared 23% amino acid sequence similarity with the closest genetic relative, *Hymenopteran flavi-related virus* (59) (Supplementary Table 2). Chalinolobus tuberculatus metagenome derived flavivirus 1 fell within the pesti-large genome flaviviruses clade infecting invertebrate hosts and thus is likely dietary-related (Figure 5b). Two additional viral transcripts, Chalinolobus tuberculatus flaviviruses 2 and 3, shared 66.2% sequence similarity to 65.91% and fell alongside members of the *Flavivirus* genera that had previously been identified in invertebrate species in Turkey and Senegal (61,62) (Figure 5b, Supplementary Table 2).

#### 4.3.3 Hepeviridae

Six novel hepeviruses were identified in the lesser short-tailed and long-tailed bat guano metatranscriptomes and were termed Mystacina tuberculata metagenome derived hepevirus 1-4 and Chalinolobus tuberculatus metagenome derived hepevirus 1-2 (Figure 5c, Supplementary Table 2). All six novel hepeviruses described shared <90% amino acid sequence similarity with their closest known genetic relatives, indicating that these viruses were novel (Supplementary Table 2). The closest genetic relatives to these viruses were to avian and reptilian metagenome-associated viruses (33,63–65) (Figure 5c). It is therefore likely that all hepeviruses identified in this study are also dietary, environmental, or commensal in origin (Figure 5c, Supplementary Table 2). A further hepevirus was discovered which shared greater than 90% amino acid sequence similarity with the closest known genetic relative, Tuatara cloaca-associated hepevirus-1. The Tuatara cloaca-associated hepevirus-1 was previously discovered in cloacal swabs from tuatara (*Sphenodon punctatus*) indicating that this virus is likely dietary, commensal or environmental in origin (33).

#### 4.3.4 Nodaviridae

We identified eight nodaviruses within the bat guano metatranscriptomes of both long-tailed bats and lesser short-tailed bats (Figure 5d). One of these viruses had previously been discovered, sharing >96% amino acid sequence similarity with *Bat nodavirus* (66). This virus had been identified in the brain of a Serotine bat but was classified as an invertebrate host-related virus, suggesting a dietary association in the context of our study too (Supplementary Table 2). The remaining seven nodaviruses were classified as novel and were provisionally named Mystacina tuberculata metagenome derived nodavirus 1-5 and Chalinolobus tuberculatus metagenome derived nodavirus 1-2. Their closest known genetic relatives were a number of vertebrate host metagenome-associated viruses (56,65–68). It is therefore highly likely that these novel nodaviruses are dietary, commensal, or environmental in origin (Supplementary Table 2).

#### 4.3.5 Peribunyaviridae

Two peribunyaviruses were uncovered in the bat guano metatranscriptomes of long-tailed bats sampled from the Grand Canyon Cave and lesser short-tailed bats sampled from Pureora Forest Park (Figure 5e, Supplementary Table 2). The novel viruses were provisionally termed Mystacina tuberculata metagenome derived peribunyavirus and Chalinolobus tuberculatus metagenome derived peribunyavirus. The two Chalinolobus tuberculatus metagenome derived peribunyavirus contigs identified in this study shared 54% and 41% amino acid sequence similarity with their closest known genetic relatives, *Hubei insect virus 1* and *Gonipterus platensis bunyan-like virus*, respectively, previously identified in arthopods and a Gum-tree weevil (*Gonipterus platensis*) (69,70), indicating that these viruses are likely dietary related (Supplementary Table 2).

A protein structural similarity based approach was used to identify the highly divergent viral contig termed Mystacina tuberculata metagenome derived peribunyavirus. The top PHYRE2 structural match was to a *La Crosse virus* RNA-directed RNA polymerase l, a member of the *Peribunyaviridae* family, modelled with a confidence of 92%, a percentage identity of 28% and a coverage of 39%. Upon analysing the amino acid sequence of Mystacina tuberculata metagenome derived peribunyavirus along with other reference *Peribunyaviridae* polymerase sequences, including the *La Crosse virus*, the Mystacina tuberculata metagenome derived peribunyavirus shared multiple conserved residues within the highly conserved RdRp motifs B-E, further supporting the view that this contig is viral (Figure 6). Upon phylogenetic analysis, Mystacina tuberculata metagenome derived peribunyavirus fell within a sister clade to *Soft tick bunyavirus*, which was previously identified in soft ticks (*Argas vespertilionis*) collected from bat guano in Japan (71), indicating that this virus may be dietary related (Figure 5e).

**Figure 6.**
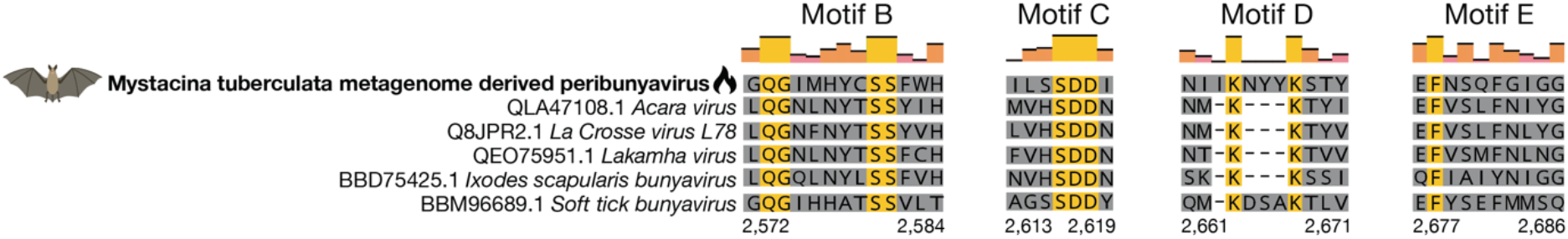
Amino acid alignment of reference *Peribunyaviridae* RdRp sequences and the novel and highly divergent Mystacina tuberculata metagenome derived peribunyavirus (bold) showing motifs B-E. The MAFFT alignment and the L-INS-I algorithm were selected to align the sequences in Geneious Prime (39). The flame symbol highlights that the Mystacina tuberculata metagenome derived peribunyavirus was identified using a protein structural similarity based approach rather than an amino acid sequence similarity based approach.

#### 4.3.6 Picobirnaviridae

Nine novel viruses within the *Picobirnaviridae* were identified in the bat guano metatranscriptomes of lesser short-tailed bats sampled from Pureora Forest Park and were termed Mystacina tuberculata metagenome derived picobirnavirus 1-4 (Figure 5f, Supplementary Table 2). The closest known genetic relatives were viruses sampled from vertebrate host metagenomes indicating that the true host of the novel Mystacina tuberculata metagenome derived picobirnaviruses 1-9 are likely dietary, environmental, or commensal in origin (56,72,73).

An additional two novel viruses were identified in the bat guano metatranscriptomes of long-tailed bats sampled from the Grand Canyon Cave and were named Chalinolobus tuberculatus metagenome derived picobirnavirus 1 and 2 (Figure 5f). Their closest genetic relative was *Limbe picobirna-like virus* (amino acid sequence similarity ∼56%) previously identified in a straw-coloured fruit bat (*Eidolon helvum*) but was classified as an invertebrate host virus (74), suggesting that the novel picobirnavirus is likely dietary related (Supplementary Table 2).

#### 4.3.7 Reovirales

We identified five novel reoviruses within the bat guano metatranscriptomes of lesser short-tailed bats sampled from Pureora Forest Park and long-tailed bats sampled from the Grand Canyon Cave (Figure 7). All five novel reoviruses termed Mystacina tuberculata metagenome derived reovirus 1-4 and Chalinolobus tuberculatus metagenome derived reovirus are likely members of the *Spinareoviridae* (Figure 7). We also uncovered a viral contig that shared 97.2% amino acid sequence similarity with *Avian associated reo-like virus 3* previously identified in the metagenome of a New Zealand fantail (*Rhipidura fuliginosa*) (Supplementary Table 2) (64).

**Figure 7.**
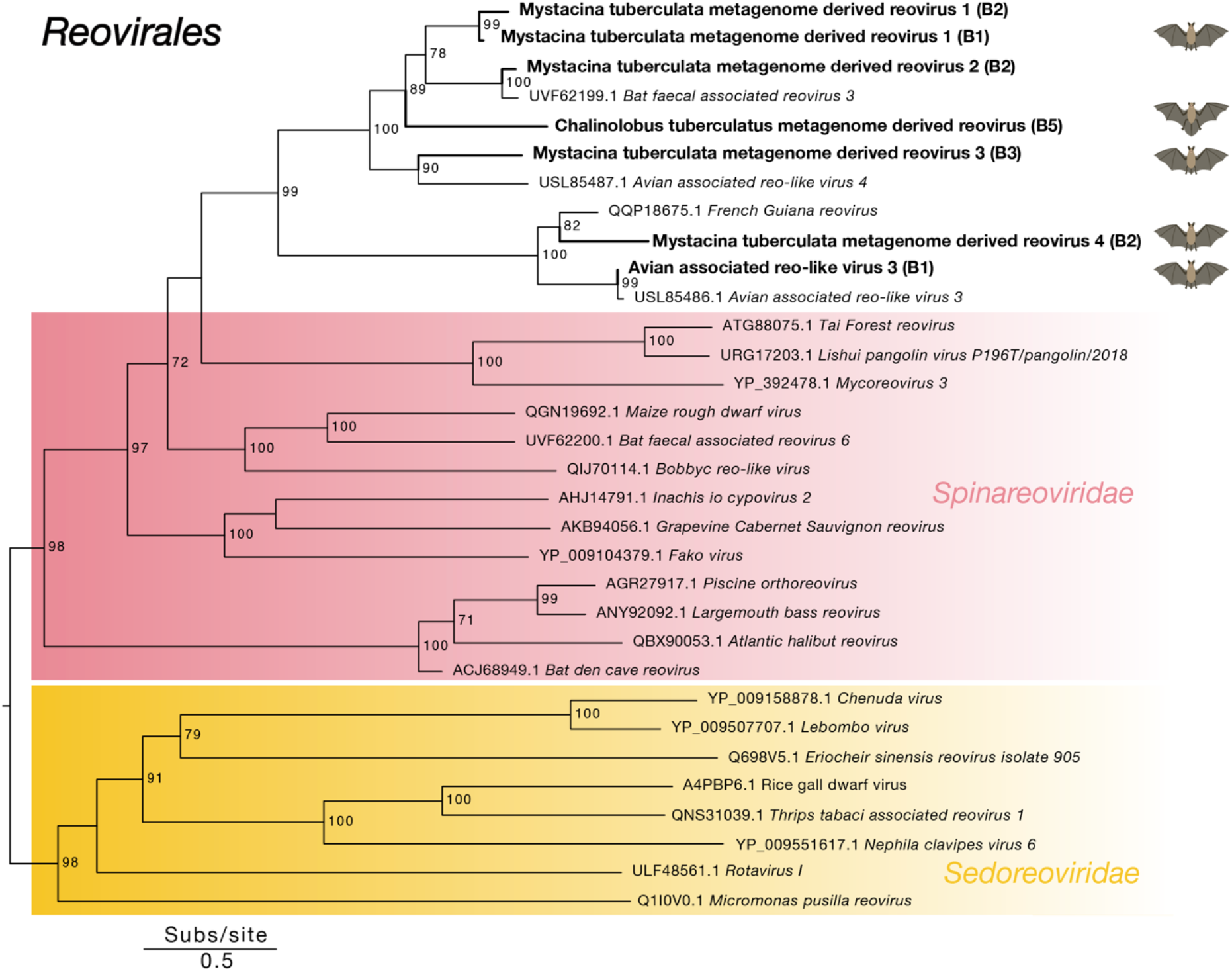
Midpoint rooted maximum likelihood phylogenetic tree of representative viral transcripts containing RdRp from the *Reovirales* order. Bat metagenome derived viruses identified in this study are in bold while viral families are highlighted. Branches are scaled to the number of amino acid substitutions per site. Nodes with ultrafast bootstrap values of >70% are noted.

### 4.4 Factors Shaping the Diversity of Bat Viromes

We next asked whether the alpha diversity of bat viromes was influenced by bat species and/or sampling location (North or South Island). This analysis revealed that neither bat species nor location significantly influenced alpha diversity of bat viromes (p = 0.2 and 0.6, respectively) (Figure 8a and 8b). Even though long-tailed bats harboured bat viruses across seven viral families (Figure 2), all but one of these viruses were found across just two libraries sampled from the Grand Canyon Cave. Therefore, the absence of bat-specific viruses within other libraries rendered this comparison statistically insignificant. Additionally, there was a weak positive yet not significant correlation between the pairwise differences in Shannon index of bat viruses across bat libraries and the pairwise distance (in kilometers) between roost sites (Pearsons correlation = 0.19, p-value = 0.13) (Figure 8c).

**Figure 8.**
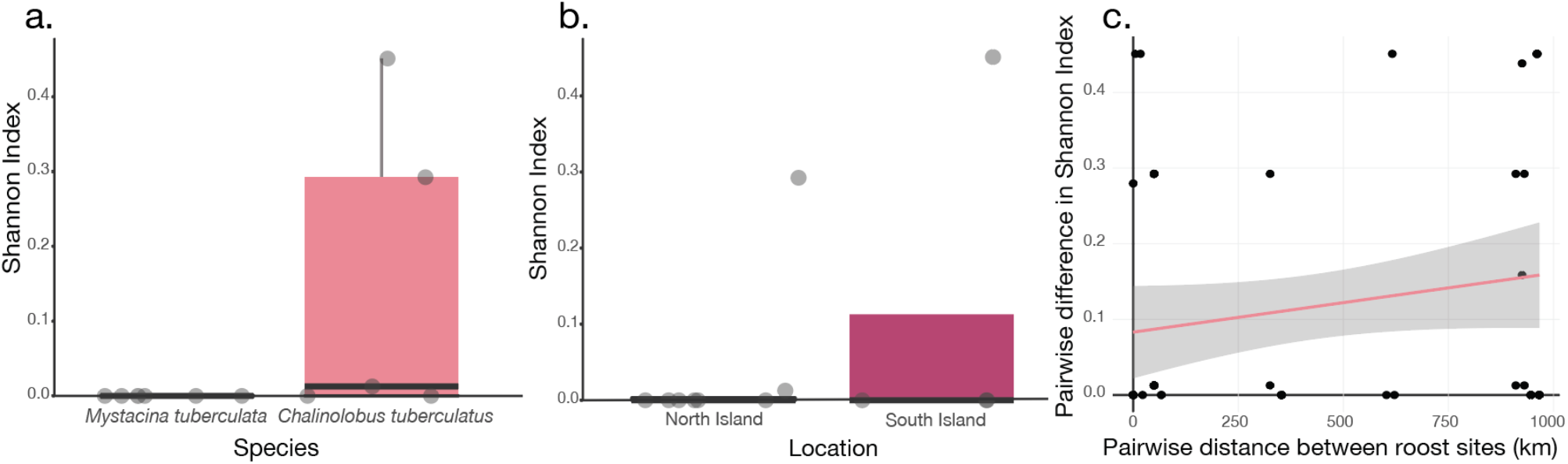
Shannon Index boxplots of (**a.**) bat viruses across bat libraries in relation to bat species and (**b.**) bat viruses across bat libraries in relation to sample location (North or South Island). Scatter plot (**c.**) of pairwise differences in Shannon index of bat viruses across bat libraries in relation to the pairwise distances between roost sites (km).

Similarly, when we considered bat viromes comprised of bat viruses as well as bat metagenome derived viruses, neither bat species nor location influenced the alpha diversity (p-value = 0.3 and 0.1, respectively) (Supplementary Figure 1a). In contrast, when considering only the metagenome derived virome, alpha diversity was significantly higher in lesser short-tailed bats compared to long-tailed bats (p-value = 0.04) (Supplementary Figure 1b), likely because lesser short-tailed bats consume a broader diet than long-tailed bats and may therefore be exposed to a wider range of dietary related viruses (12,27). Similarly, bats that were sampled from the North Island were significantly more diverse in their metagenome derived viromes in comparison to bats that were sampled from the South Island (p-value = 0.02) (Supplementary Figure 1b). However, the majority of samples from the North Island were from lesser short-tailed bats (four out of six) in comparison to only one (out of four) sampled from the South Island.

## 5. Discussion

We analysed the viromes of Aotearoa New Zealand’s only native terrestrial mammalian species, the lesser short-tailed bat and the long-tailed bat, to determine patterns of cross-species virus transmission and whether their contrasting host ecologies and life histories have influenced their virome composition. Our analysis resulted in the discovery of many bat viruses in guano samples from long-tailed bats, yet no *bona fide* bat viruses in lesser short-tailed bats were identified. At face value this suggests that long-tailed bats may be a more important reservoir compared to lesser short-tailed bats. Notably, viruses spanning the families *Picornaviridae*, *Astroviridae*, *Paramyxoviridae*, *Papillomaviridae*, *Poxviridae*, *Rhabdoviridae* and *Coronaviridae* were identified in guano metatranscriptomes of long-tailed bats, including, to our knowledge, the shortest known coronavirus genome documented to date. These novel bat viruses were all closely related to others identified in bats. Prior to this study, only short transcripts from two viruses, both alphacoronaviruses, had been identified in long-tailed bat guano (17). Consequently, this study has expanded our knowledge of the viruses that circulate within New Zealand’s long-tailed bat populations.

The Grand Canyon Cave, home to a population of long-tailed bats located in the central North Island of New Zealand, was where all but one of the novel bat viruses were discovered. Long-tailed bats often move between roosts and primarily roost during the day in small colonies distributed among multiple tree-roosts in native forests (75), yet are also known to roost at night in large numbers in the Grand Canyon Cave, particularly during spring (12,76). The maximum number of night roosting long-tailed bats that were counted during any one time in the Grand Canyon Cave was 358, while a total of 533 individuals were banded over three seasons indicating that bats from multiple different day roosting colonies may come together to roost in the cave at night (76). While the reason behind night roosting in long-tailed bats is unclear (76), the Grand Canyon Cave likely provides an ideal site to facilitate viral transmission and thus high viral richness as demonstrated here. Examining the role that bat roosting behaviour plays in the transmission of viruses within New Zealand’s bat species would add to a better understanding of the potential risk of viral spillover.

In previous studies, short virus transcripts from the *Coronaviridae*, *Papillomaviridae*, *Picornaviridae*, *Polyomaviridae*, *Caliciviridae* and *Hepeviridae* have been uncovered in guano from lesser short-tailed bats located on Whenua Hou (Codfish Island), a small (14 km^2^), predator-free, offshore island in southern New Zealand (15,16). Whenua Hou is home to a natural population of more than 2000 short-tailed bats that occupy more than 120 tree roosting sites across the island (77,78). While we also identified novel viral contigs belonging to the *Hepeviridae* in this host, phylogenetic analysis placed these contigs as most likely to be dietary, commensal, or environmental in origin as they were closely related to viruses identified in the metagenomes of avian and reptilian species. Here, we were unable to detect any bat viruses in guano sampled from mainland populations of lesser short-tailed bats.

We identified two alphacoronaviruses in long-tailed bat guano that were closely related to alphacoronaviruses previously identified in lesser short-tailed bats. Although there is no evidence that lesser short-tailed bats and long-tailed bats occupy the same roosts (29), roost sites can be within close proximity where both species may interact (12,77). That both bat species harbour the same viral species, such that these viruses have jumped between these hosts, provides evidence of this interaction. The large geographical distance between where these viruses where sampled (>1000 km) indicates that these viruses may be present across multiple roosts throughout New Zealand, although we were unable to detect them across other locations sampled here. Longitudinal sampling of bats across New Zealand may shed new light on the persistence of viruses within these populations, particularly as the rate of viral shedding amongst bats is known to vary throughout the year (79).

Long-tailed and lesser short-tailed bats are highly divergent, belonging to different families. As such, the incongruence observed between viral and host topologies supports the cross-species transmission of coronaviruses. Both the long-tailed bat alphacoronaviruses identified here were closely related to those present in Gould’s wattled bat from Australia (26). Like New Zealand’s long-tailed bats, these bats are members of the Vespertilionidae. It is likely, then, that long-tailed bats were reservoirs of coronaviruses before migrating to New Zealand and transmitted these viruses to lesser short-tailed bats since their arrival (12,22). This new evidence suggests that estimates of the time to most recent common ancestor of orthocoronaviruses may require further calibration (47), although such analyses are challenged by uncertainty over the timing of bat migrations to New Zealand (∼1-17mya for long-tailed bats and ∼16-35mya for lesser short-tailed bats) estimated from fossil records and species divergence times (12,22–25). Clearly, obtaining full-length genomes of alphacoronaviruses previously identified in lesser short-tailed bats is an important step to arriving at a more comprehensive understanding of their evolutionary history.

While coronaviruses typically range from 26-32 kilobases in length (80), we identified an alphacoronavirus of only 22,416 nucleotides, containing a large deletion in ORF1a of both NSP1 and NSP2, as well as a truncated NSP3. NSP1 is involved in inhibiting the host immune response, decreasing the overall expression of genes in the host cells while redirecting host machinery to produce viral proteins (81). While gammacoronaviruses and deltacoronaviruses are known to lack NSP1, all currently known alpha and betacoronaviruses contain NSP1 (81–83). The function of NSP2 is currently unclear (84), although deletion of NSP2 in two other betacoronaviruses –*Severe acute respiratory syndrome coronavirus 1* and *Murine hepatitis virus* – still resulted in viable viruses in cell culture, although viral growth and RNA synthesis was reduced (85). NSP3 is involved in a number of roles including polyprotein processing, acting as a scaffold protein and is an essential component of the replication and transcription complex (86). To the best of our knowledge this alphacoronavirus may be the shortest coronavirus genome identified to date. Despite its sequence similarity within the RdRp, it’s shortened genome and divergence in other genes suggests that it should be classified as a novel virus species.

Two novel viruses from the *Flaviviridae* were uncovered in the guano of both bat species. Flaviviruses have been known to infect a wide range of hosts and cause disease in humans, livestock and wildlife (87). Both novel viruses identified here fell within the ‘pesti-large genome’ clade (60). The pesti-large genome clade is associated with invertebrate hosts (60), making it likely that these viruses are dietary in origin (12,27). One virus, Pekapeka metagenome derived flavivirus, was especially divergent, expanding the diversity of this clade and highlighting the potential to uncover highly divergent novel viruses within hosts that have been severely understudied.

In sum, this study has increased our understanding of the viruses that circulate within New Zealand’s bat populations and indicates that coronaviruses have likely jumped between these bat populations since long-tailed bats arrived at least two million years ago. A future focus on understanding the influence of roosting behaviour and seasonality on virome composition and further sampling New Zealand’s mammalian hosts will be important to expand our knowledge of viral diversity in New Zealand bats and determine whether the bat viruses documented are present in other terrestrial mammalian hosts.

## 8. Acknowledgements

We would like to thank the Department of Conservation for coordinating the sampling of the bat guano from multiple sites across New Zealand. We would also like to thank local iwi for supporting our research.

## 9. Supplementary Material

**Supplementary Table 1**. Data surrounding bat library pooling and sampling site locations.

**Supplementary Table 2**. Data surrounding the viral contigs that were identified in this study and the top BLASTp results.

**Supplementary Figure 1**. Shannon index analysis of (**a.**) bat viruses and bat metagenome derived viruses across bat libraries in relation to bat species (left), sample location (North or South Island) (center) and the pairwise distance between roost sites (km) (right). Shannon index analysis of (**b.**) bat metagenome derived viruses across bat libraries in relation to bat species (left), sample location (North or South Island) (center) and the pairwise distance between roost sites (km) (right).

## 10. Data Availability

The raw sequencing reads generated in this project are available on the Aotearoa Genomic Data Repository, DOI: (pending); while the virus sequences are available under GenBank accessions (pending) (Supplementary table S2). Alignments and code for the statistical analysis can be found at https://github.com/stephwaller/NZ-Bat-Virome-Paper.git.

