## Supplementary figures and images for "Virome analysis of New Zealand’s bats reveals cross-species viral transmission among the *Coronaviridae*"

### Supplementary Figure 1

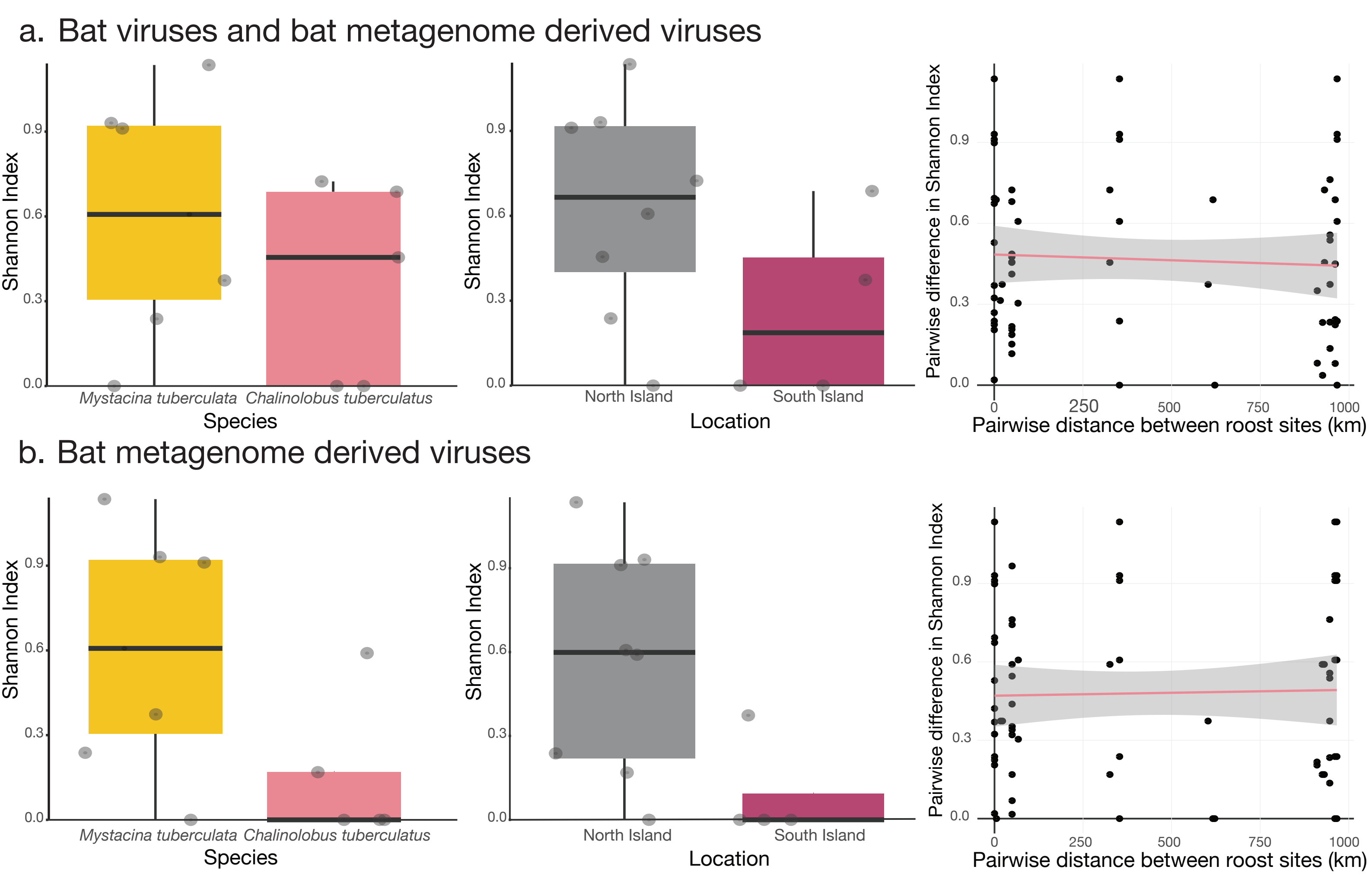
